# Quantifying and understanding well-to-well contamination in microbiome research

**DOI:** 10.1101/577718

**Authors:** Jeremiah J Minich, Jon G Sanders, Amnon Amir, Greg Humphrey, Jack Gilbert, Rob Knight

## Abstract

Microbial sequences inferred as belonging to one sample may not have originated from that sample. Such contamination may arise from laboratory or reagent sources or from physical exchange between samples. This study seeks to rigorously assess the behavior of this often-neglected between-sample contamination. Using unique bacteria each assigned a particular well in a plate, we assess the frequency at which sequences from each source appears in other wells. We evaluate the effects of different DNA extraction methods performed in two labs using a consistent plate layout including blanks, low biomass, and high biomass samples. Well-to-well contamination occurred primarily during DNA extraction, and to a lesser extent in library preparation, while barcode leakage was negligible. Labs differed in the levels of contamination. DNA extraction methods differed in their occurrences and levels of well-to-well contamination, with robotic methods having more well-to-well contamination while manual methods having higher background contaminants. Well-to-well contamination was observed to occur primarily in neighboring samples, with rare events up to 10 wells apart. The effect of well-to-well was greatest in samples with lower biomass, and negatively impacted metrics of alpha and beta diversity. Our work emphasizes that sample contamination is a combination of crosstalk from nearby wells and background contaminants. To reduce well-to-well effects, samples should be randomized across plates, and samples of similar biomass processed together. Researchers should evaluate well-to-well contamination in study design and avoid removal of taxa or OTUs appearing in negative controls, as many will be microbes from other samples rather than reagent contaminants.

**Importance:** Microbiome research has uncovered magnificent biological and chemical stories across nearly all areas of life science, at times creating controversy when findings reveal fantastic descriptions of microbes living and even thriving in once thought to be sterile environments. Scientists have refuted many of these claims because of contamination, which has led to robust requirements including use of controls for validating accurate portrayals of microbial communities. In this study, we describe a previously undocumented form of contamination, well-to-well contamination and show that contamination primarily occurs during DNA extraction rather than PCR, is highest in plate-based methods as compared to single tube extraction, and occurs in higher frequency in low biomass samples. This finding has profound importance on the field as many current techniques to ‘decontaminate’ a dataset simply relies on an assumption that microbial reads found in blanks are contaminants from ‘outside’ namely the reagents or consumables.

## Introduction

Massively high-throughput sequencing has enabled fundamental changes to the study of microbial ecology. Increased throughput and sequencing depth has empowered researchers to utilize multiplexing to increase sample sizes to thousands per study [1–6]. However, new ways of knowing require new understanding of potential flaws and confounds. Many studies have addressed computational and statistical challenges associated with analyzing 16S rRNA gene sequence data, including the impacts of sequence similarity clustering [7], diversity estimation, and data compositionality [8], to name just a few. There has also been substantial effort to reduce confounding experimental effects via standardization of microbiome sample processing methods, including sample collection, preservation [9], DNA extraction [10–12], library preparation [6,13–17] and sequencing [5]. Together, these approaches have facilitated large-scale metaanalyses such as the Earth Microbiome Project ‘EMP’ (earthmicrobiome.org) [2]. Despite these efforts, a significant amount of experimental noise remains in any given microbiome study.

Contamination, or the observation of sequence reads in a sample coming from microbes that weren’t originally part of that sample, remains one of the most pernicious types of experimental noise. Microbial rRNA gene copies can be found even in ‘sterile’ reagents, leading to that presence of background signal derived from DNA extraction kits [18], PCR mastermix [19], and other consumables [20]. It is now widely understood that such contaminants must be considered in microbiome analyses, especially when dealing with low-biomass samples where contaminant rRNA gene copies make up a larger fraction of the community [7,21–25]. Various engineering strategies have been proposed and are utilized to minimize contamination including physical separation of rooms used for DNA extractions and PCR, wearing additional PPE [26] to cover skin to prevent technician-induced contaminants, UV sterilization of plastic consumables or reagents, or ethidium oxide treatment of consumables [20].

Beyond physically limiting contamination, the use of positive and negative controls is increasingly being used to assess and quantify contamination in a study, allowing for the potential of contaminant removal in silico [27]. Methods such as Katharoseq [11], utilize the ratio of read counts and composition of positive and negative controls, to determine criteria for sample inclusion. Others have emphasized the importance of including negative controls to understand background contamination [28]. Based on the idea that contaminants are primarily derived from external sources, some have proposed the strategy of simply identifying this ‘contaminome’ profile and then removing them from the dataset [29]. This, however fails to contend with the potential that contaminants may arise from other samples within a study itself. Such between-sample contamination has been observed as a product of ‘barcode-swapping’ between samples as a byproduct of Illumina ex-Amp sequencing reactions, and has also been suggested to arise from improper assignment of barcodes to neighboring clusters in image processing [30]. Anecdotally, we have also observed instances that appear to arise from physical cross-contamination of samples. Since most DNA extractions and PCR reactions are performed on multiple samples at once, often times in 96-well format, we reasoned it would be important to take into consideration that nearby samples could in fact contribute to contamination of negative controls.

To evaluate this hidden factor of contamination, we designed an experiment to empirically characterize the frequency and nature of well-to-well contamination using different DNA extraction and sample handling protocols. By placing 16 unique bacterial “source” isolates at high biomass in individual wells across plates of alternating low-biomass “sink” bacteria and no-template blank wells, we were able to observe and quantify well-to-well transfer events under different scenarios, including automated plate-based extraction and manual tube-based extraction protocols. We further included libraries from an additional, unique, isolate that were extracted and amplified separately to account for potential instrument-based cross-contamination mechanisms such as barcode-swapping or miss-assignment. To further validate results, we processed an additional two 96-well plates at another microbiome facility.

## Results

We designed a 96-well plate layout containing 16 unique source bacteria (~10,000,000 cells per well, corresponding to 10^8^ cells ml^−1^), 24 sink wells (containing V. fischeri at ~100,000 cells per well, 10^6^ cells ml^−1^) and 48 blank wells (Figure 1a). At UCSD, a total of three replicate sample plates were DNA extracted: two using the Epmotion5075 with magnetic bead cleanups on Kingfisher robots (Plate 1, Plate 2) and one manually with column cleanups (Tube). All three plates were then processed each with two unique PCR plates in triplicate as outlined (Figure 1). In addition, 16 gDNA replicates of a *Clostridium* isolate were processed on its own 96-well plate and amplified in a separate PCR reaction, to allow for detection of instrument-based barcode miss-assignment. A mock community comprised of all source isolates and the sink isolate was created and then serially diluted and processed as well to validate sample amplification. Details on the actual plate map patterns can be found in Additional file 1 for all eight PCR plates. A total of 3,756,064 reads from 713 samples resulted in 6305 features. A summary table was generated to describe well-to-well and background contamination occurrences across the samples (Additional Table 1). One of the 16 source microbes was highly contaminated with background contaminants and did not produce the expected sequence results, but was included in the analysis as we did not want to bias our results.

**Table 1:** Summary of contamination (well to well and background) impact on NTCs, low biomass, and high biomass sample types

|  |  |  | (Well-to-well) |  |  |  |  |  | (background-kits) |  |  |
| --- | --- | --- | --- | --- | --- | --- | --- | --- | --- | --- | --- |
|  |  |  | Prev | Richness |  | Composition |  |  | Composition |  |  |
| Type | Location | Extract | mean | Total | W2W | mean | median | max | mean | median | max |
| ntc(n) |  |  |  |  |  |  |  |  |  |  |  |
| 61 | UCSD | m_tube | 0.4754 | 20 | 0.041 | 0.046 | 0 | 0.56 | 0.9382 | 1 | 1 |
| 32 | Argonne | m_tube | 0.5313 | 165 | 0.016 | 0.009 | 0.0003 | 0.0823 | 0.9915 | 0.9997 | 1 |
| 28 | Argonne | m_plate | 0.1071 | 8 | 0.042 | 0.031 | 0 | 0.7517 | 0.9686 | 1 | 1 |
| 116 | UCSD | r_plate | 0.9569 | 15 | 0.278 | 0.583 | 0.6579 | 1 | 0.3307 | 0.2025 | 1 |
| sink(n) |  |  |  |  |  |  |  |  |  |  |  |
| 93 | UCSD | m_tube | 0.1505 | 20 | 0.01 | 5E-04 | 0 | 0.0278 | 0.0335 | 0.0168 | 0.9873 |
| 48 | Argonne | m_tube | 0.5 | 189 | 0.017 | 0.023 | 0 | 0.5934 | 0.7808 | 0.8382 | 0.9878 |
| 46 | Argonne | m_plate | 0.3261 | 16 | 0.066 | 0.14 | 0 | 0.9871 | 0.5846 | 0.6267 | 1 |
| 187 | UCSD | r_plate | 0.6738 | 15 | 0.127 | 0.007 | 0.0008 | 0.1561 | 0.0091 | 0.0023 | 0.4051 |
| source(n) |  |  |  |  |  |  |  |  |  |  |  |
| 31 | UCSD | m_tube | 0.6129 | 18 | 0.065 | 0.001 | 0.0001 | 0.0299 | 0.083 | 0.0029 | 1 |
| 16 | Argonne | m_tube | 0.875 | 21 | 0.138 | 2E-04 | 0.0002 | 0.0007 | 0.1154 | 0.0041 | 0.9998 |
| 16 | Argonne | m_plate | 0.8125 | 17 | 0.168 | 0.024 | 0.0001 | 0.364 | 0.1313 | 0.0032 | 0.9999 |
| 64 | UCSD | r_plate | 0.7031 | 12 | 0.138 | 0.009 | 0.0004 | 0.5067 | 0.0732 | 0.0016 | 1 |
| Prev-prevalence (number of samples with any well-to-well contamination / total number of samples) |  |  |  |  |  |  |  |  |  |  |  |
| Extract: DNA extraction method used |  |  |  |  |  |  |  |  |  |  |  |
| m-manual (non-robotic based extraction) |  |  |  |  |  |  |  |  |  |  |  |
| r-robot based DNA cleanup |  |  |  |  |  |  |  |  |  |  |  |

**Fig 1:**
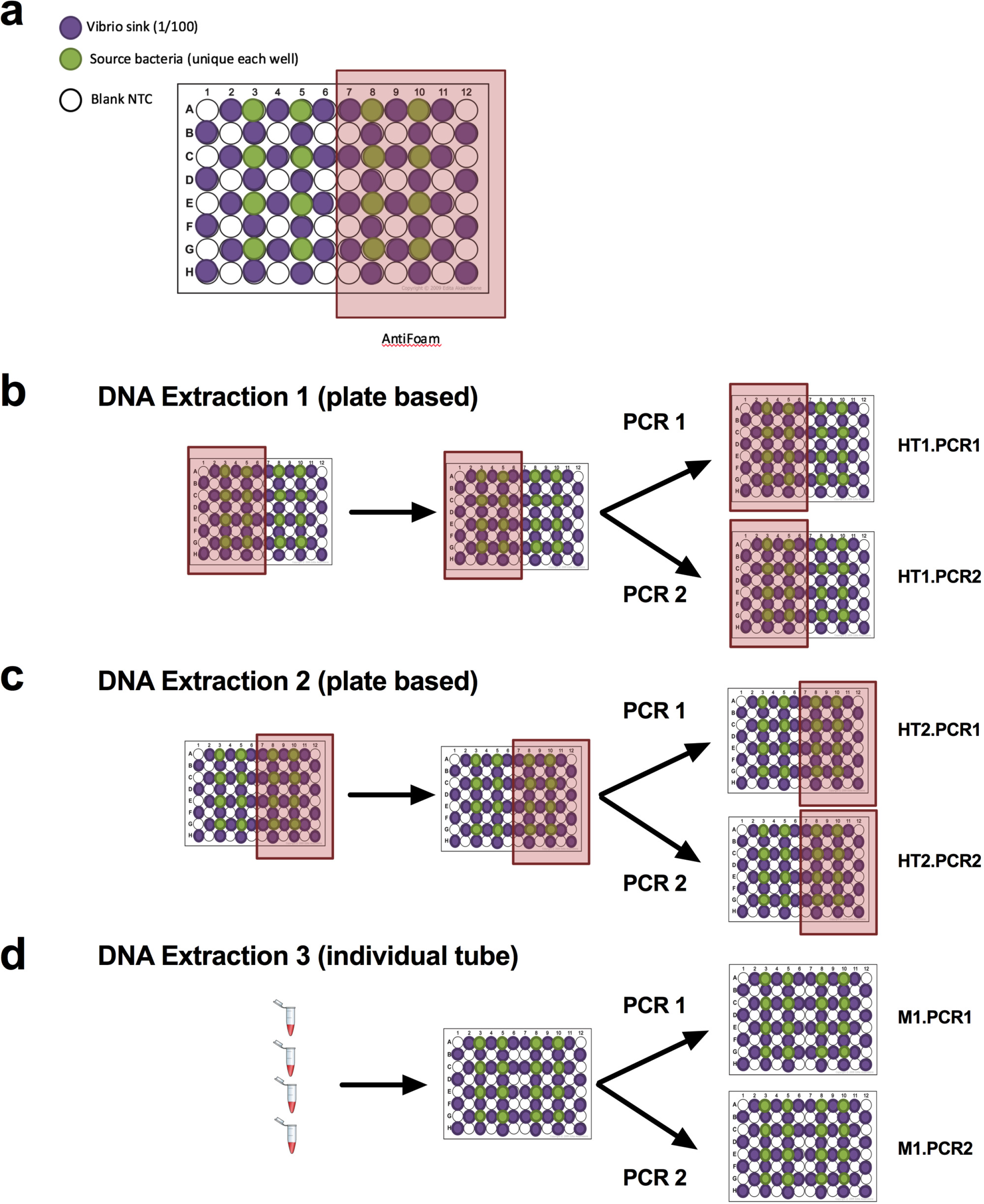
Plate design, experimental design. (a) ntc ‘white’, sink ‘purple’, and source ‘green’ samples are distributed in a checkboard pattern across the plate. Antifoam A is added to last half (b) and first half (c) of the 96-well plates processed with the robot. The manual samples did not get antifoam A. Each unique DNA extraction plate is processed in duplicate PCR plates.

Well-to well contamination events were analyzed by counting fraction of reads from a given source well appearing in other source wells, low biomass sink wells, or blanks. In our setup, well-to-well contamination was visualized to occur in all six PCR replicate plates in both labs. Based on the visualized plate patterns, the pattern of well-to-well contamination was observed to be higher in plate extractions compared to tube extractions, and was more prominent in wells directly surrounding the source well, suggesting a physical mechanism for well-to-well contamination (Figure 2). We quantified the distance by measuring contamination counts as a function of the Pythagorean distance from the source well, and determined that the highest rates of contamination occurred in the immediate proximate wells for both plate and tube extractions, but with a stronger distance-decay relationship for the plate vs. the tube extractions (Figure 3). The supplementation of Antifoam-A to wells during DNA extraction did not reduce well-to-well contamination (Additional file 2).

**Fig 2:**
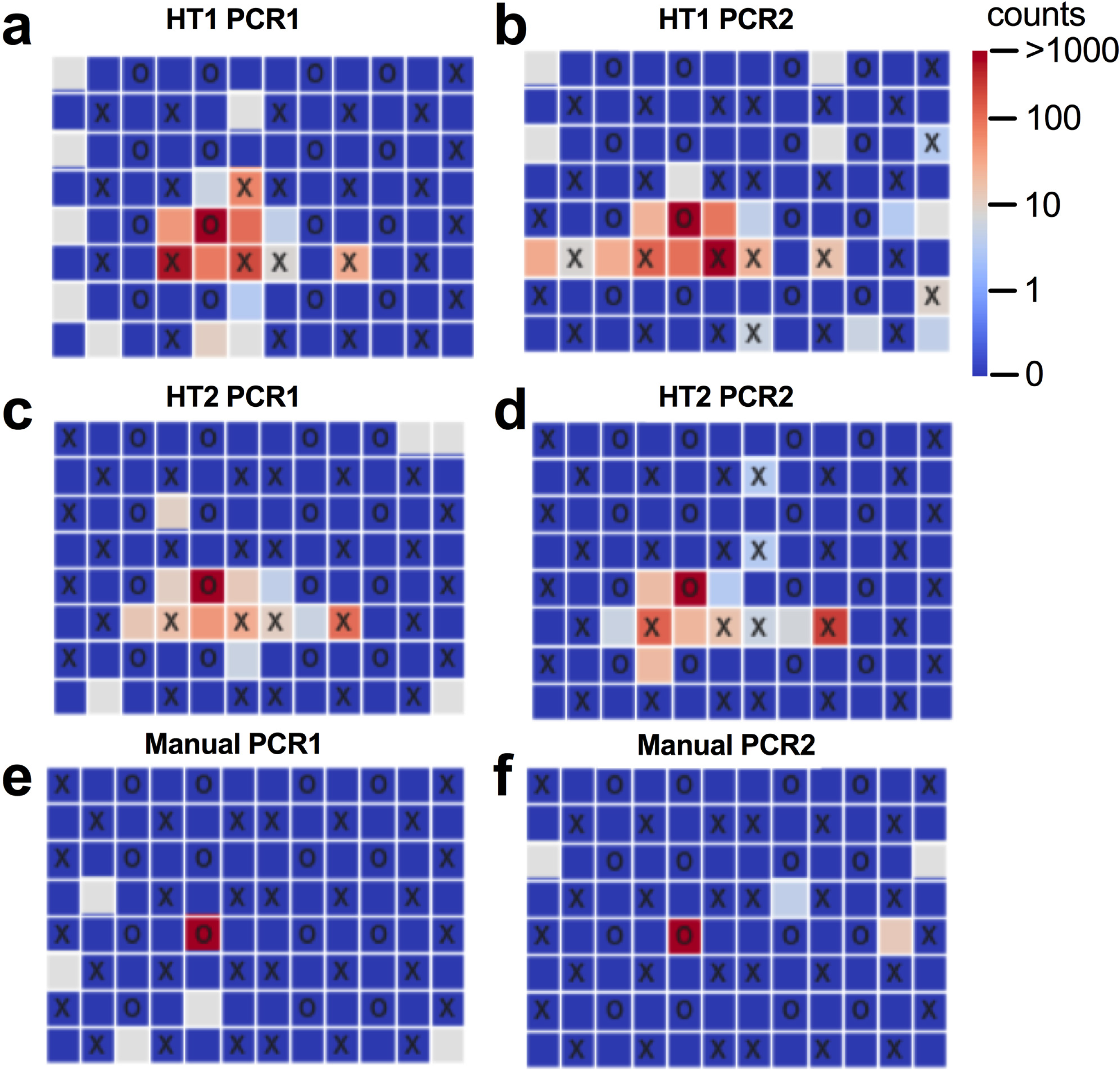
Example of plates with cross contamination; Each panel depicts a 96-well plate with source, sink and blank wells denoted by “X”, “O” empty squares respectively. Colors indicate the number of reads from a specific bacteria (*Psychrobacter spp.*, present in well E5). Panels a,b c,d and e,f correspond to two PCR replicates of robotic extraction 1,2 and manual extraction respectively.

**Fig 3:**
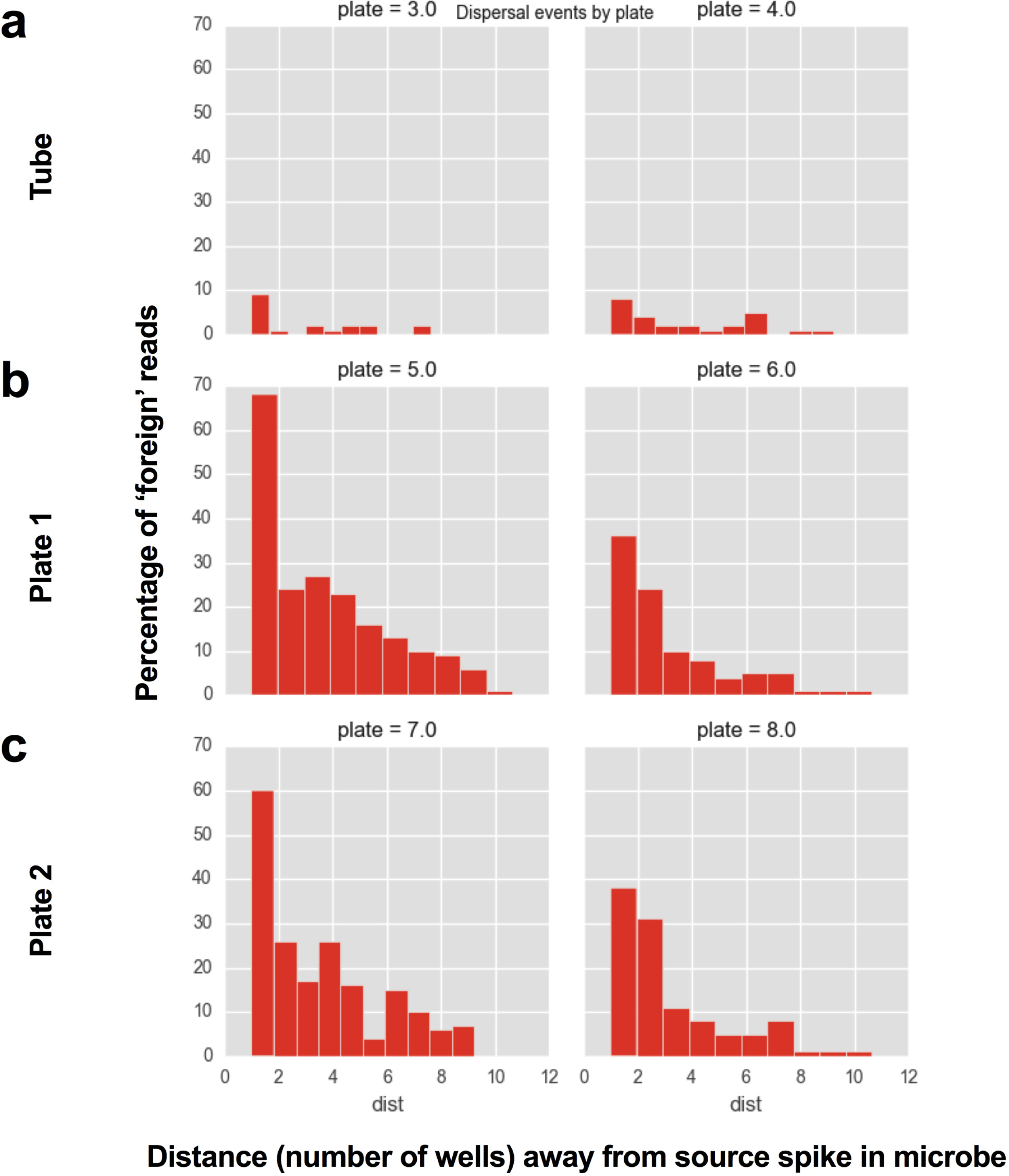
Distance decay relationship

Another possible contributing source of inter-sample contamination is barcode leakage, i.e. reads originating from a given sample being identified as originating from a different sample due to read errors in the barcode. Such “barcode-hopping” behavior has been observed in labs using 8 bp barcodes in the Microbiome Quality Control project [31]. In order to quantify the contribution of such events in our 12 bp barcode design, we designed another plate containing 16 replicate wells of a single *Clostridium* isolate. Since these samples were sequenced together with the PCR replicate plates, barcode leakage would be expected to results in Clostridium reads appearing in the PCR replicate plates samples. Barcode leakage was quantified by counting the number of reads originating from barcodes not present in the plate, and no such reads were observed, indicating that for the 12 bp Golay error correcting barcodes sequenced in these conditions, this is a very rare event (less than 1 in 3.75E6 reads), and is not a contributing factor to inter-sample contamination.

To further quantify the total effect of well-to-well contamination, we compared the proportion of microbial community source for each sample across the three DNA extraction plates (plate 1, plate 2, and tube) and for each of the two PCR replicate plates (PCRA and PCRB) from each extraction (Figure 4). Contamination frequency and relative abundance was highest in plate 1 followed by plate 2 and lowest in the tube plate (Additional file 3). NTCs were composed of primarily background contaminants in the tube extractions for both PCR replicates (median fraction of well-to-well reads 0). However, in some plate extraction NTCs, the majority of reads originated from well-to-well reads (median fraction of well-to-well reads of 0.78, 0.9, 0.44 and 0.77 for plate 1: PCRA, PCRB; plate 2: PCRA, PCRB respectively) (Additional file 4). Sink wells were also partially contaminated with source microbes, particularly in the Plate 1 replicate. The total occurrence (prevalence) of well-to-well contamination across the various sample types and extraction methods along with summarizing compositional effects of well-to-well contaminants on samples (mean, median and max) is detailed in Additional Table 2. For NTCs, 47.5% of blanks from tubes and 95.7% of blanks from plate extractions had well-to-well contamination. For low biomass samples, 15.0% of sink wells from tubes and 67.4% of sink wells from plate extractions had well-to-well contamination (Table 1).

**Fig 4:**
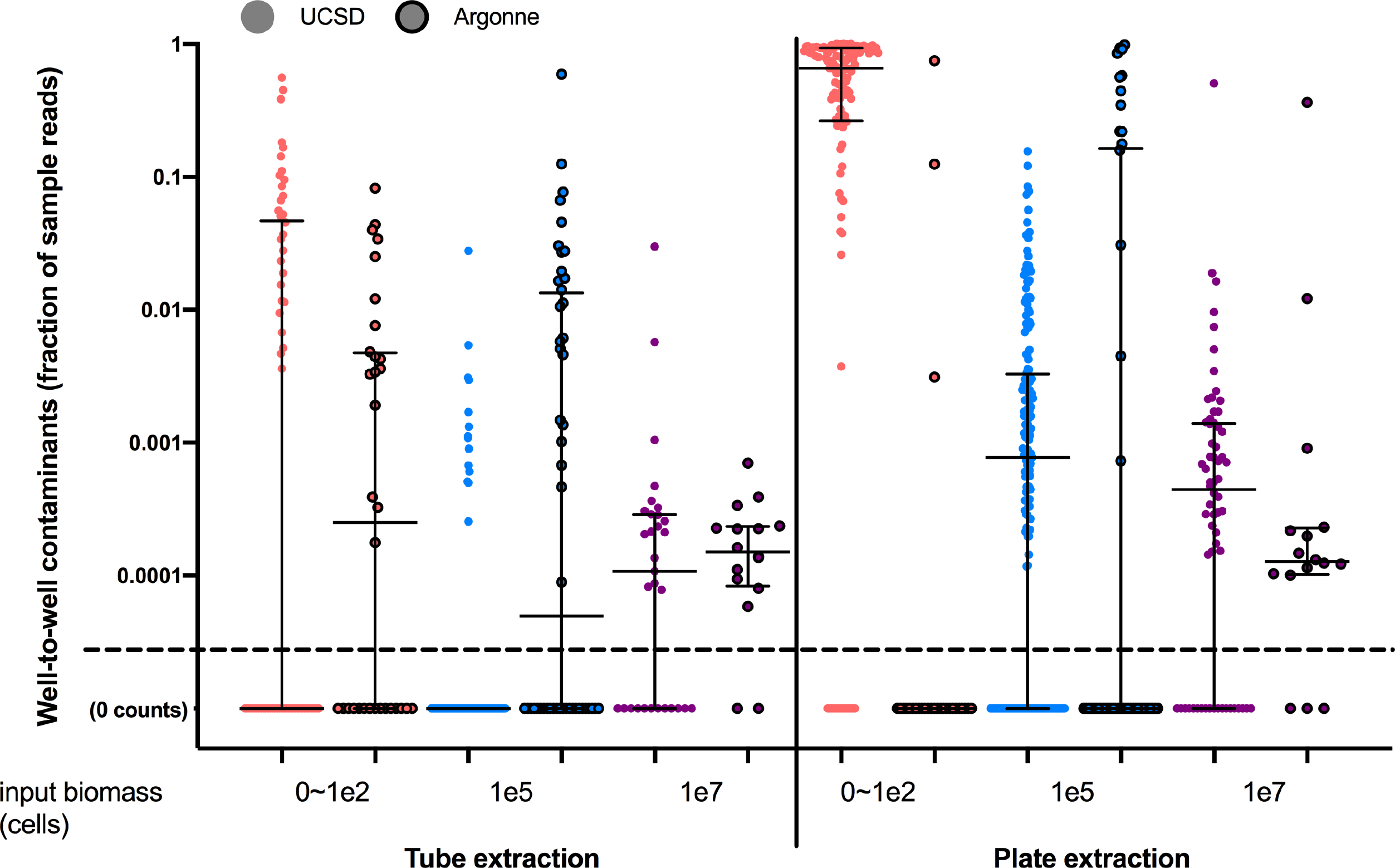
Summary statistics of sample fraction composition of well-to-well contaminants compared across extraction types (blanks - pink, sink - blue, source - purple) and across extraction methods (tube vs. plate). Samples processed at UCSD in circles with no outline and samples processed at Argonne as circles with dark border. All samples with 0 well-to-well contamination occurrences are given a count of 0.00001 to enable visualization on graph (labeled 0 counts). Median and interquartile range are displayed in black lines over the data points.

To determine if DNA extraction method (tube vs plate) had an impact on well-to-well contamination, we compared relative abundances of well-to-well contaminants for NTCs, sink, and source samples independently (Figure 5a). Well-to-well contamination was affected by extraction method, and was generally higher in plate-based extractions compared to manual single tube extractions (Kruskal-Wallis P<0.0001, Figure 5a).

**Fig 5:**
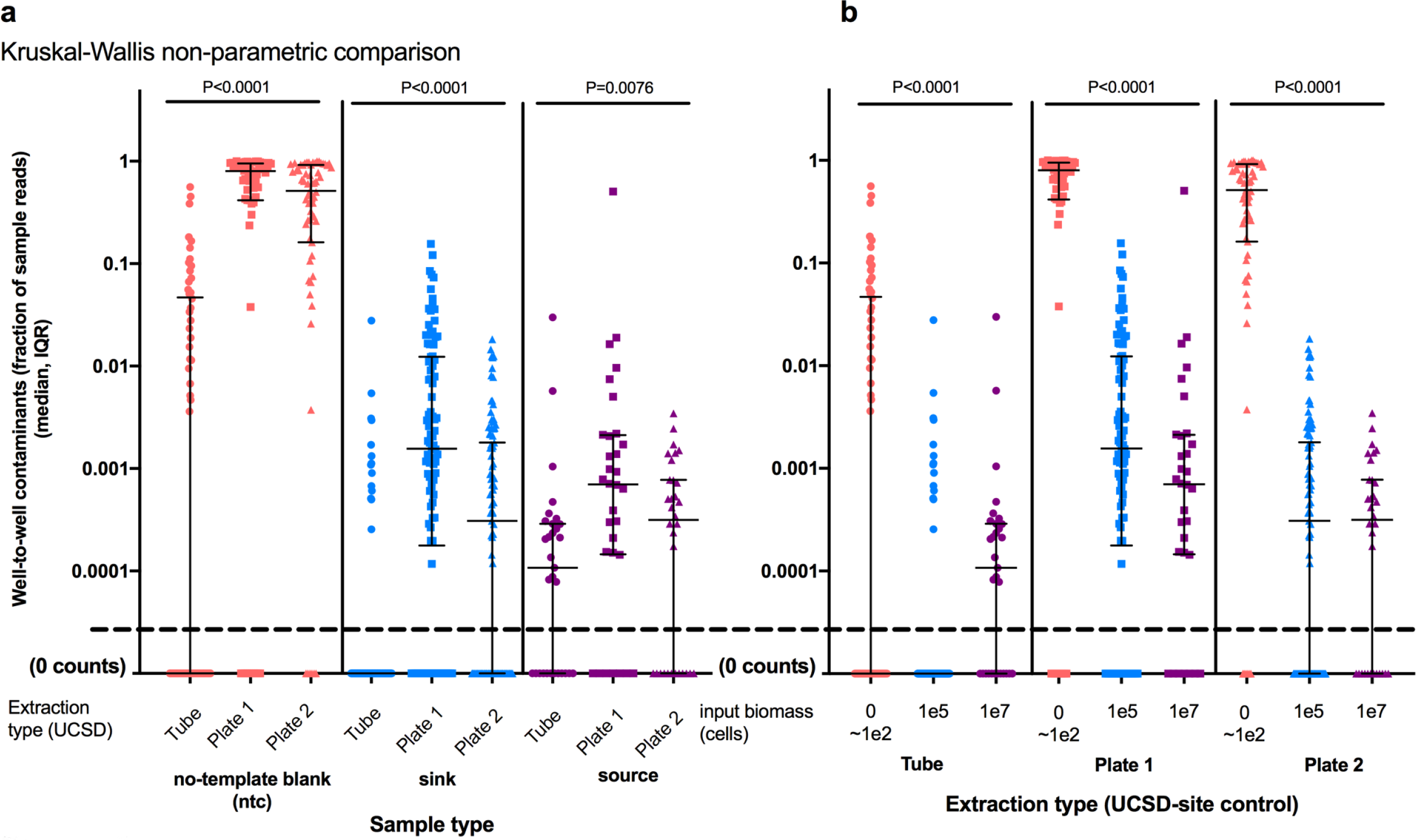
Well-to-well effect size. Proportion of sample containing well-to-well contaminants as organized by (a) sample type (ntc, sink, source) and (b) extraction method. Statistical analysis within bars performed using Kruskal-Wallis non-parametric testing.

Further, the proportion of well-to-well contamination was greater in samples with lower starting biomass (NTCs, 0-100 cells and sinks, approximately 100,000 cells) than in source wells, which had higher starting biomass (approximately 10,000,000 cells) while controlling for extraction method (Figure 5b). Well-to-well contamination was greatest in samples with lower microbial biomass.

In order to validate these results in an independent lab, in addition to the samples processed at UCSD, we sent away bacterial samples to be processed at an outside facility using the manual single tube extraction and plate extraction (although due to available facilities both utilized a column cleanup step rather than magnetic beads). All results for replicate PCR plates and robot extraction replication were summarized for overall comparison purposes (Table 1). While controlling for site (UCSD only), the total fraction of reads from samples (mean, median, and max out of 100%) caused by well-to-well contamination was highest in NTCs followed by sink and lastly source microbes for both the tube (NTC: 4.57%, 0%, 56.0%; sink: 0.05%, 0.0%, 2.78%; source: 0.13%, 0.01%, 2.99%) and plate (NTC: 58.26%, 65.79%, 100.0%; sink: 6.9%, 0.078%, 15.61%; source: 0.94%, 0.04%, 50.67%) extraction methods (Table 1 and Figure 4). The NTCs of samples processed outside of UCSD had well-to-well contamination consistent with the other tube methods while the sink samples had higher well-to-well contamination and overall background contamination than both tube and plate processed samples at UCSD (Table 1).

Since well-to-well contamination can introduce additional bacteria to samples, it has the potential to inflate alpha and decrease resolution in beta diversity metrics, especially for binary metrics (such as number of observed species, Jaccard dissimilarity, or unweighted UniFrac distance). While all of our source and sink control samples should have only had one unique sOTU, richness was typically much higher than this due to contamination including background kit contaminants along with well-to-well contaminants. We calculated the total richness per sample, which should have been one, and determined the percentage of that richness which was due to well-to-well contamination. Both well-to-well contaminants and background kit contaminants contribute to this inflated richness. Controlling for site (UCSD only), we determined that well-to-well contamination inflated richness estimates for both tube and plate extracted samples by contributing to on average (0.96%, 12.7%; tube, plate) of sink sample richness and (6.51%, 13.76%; tube, plate) of source sample richness.

We next assessed the impact of well-to-well contamination on beta-diversity measurements of the communities. Specifically, for each unique DNA extraction plate, we performed pairwise well_ID comparisons of the PCR replicates for each of the three sample types including NTCs, sink, and source microbes. Because well-to-well contamination generally only made up a small proportion of the total reads of each sample, binary metrics (which tend to emphasize the impact of rare taxa) were more affected than abundance-weighted metrics. (Additional file 5).

To further elaborate on this observation and quantify where well-to-well contamination was coming from (PCR process only or DNA extraction), we compared replicate plates which were processed using the robot. This included two separate DNA extraction plates and then two PCR plates for each extraction plate. For each PCR replicate plate, 96 pairwise distances were computed and categorized by sample type for each of the two DNA extraction plates (light red shade Additional file 5b). In addition, the pairwise distances from each of the 96 wells of the two replicate DNA extraction plates processed on the robots were also compared for the PCR replicate plate PCRA only. We found much less between-PCR than between-extraction variance, indicating that the combination of stochastic effects plus well-to-well contamination for DNA extraction is greater than for stochastic effects plus well-to-well for PCR (Additional file 5b).

## Discussion

Understanding experimental biases or noise in microbiome research is critical to drawing accurate inferences of the microbial world. Since microbes are everywhere [2], it is extremely important to limit and ideally eliminate false positives in sample signatures. Contamination is a combination of background contaminants (DNA extraction kits, PCR mastermixes, and enzymes), processing contaminants (equipment, air, technicians), and plate contaminants (well-to-well contamination). In this study, we showed that well-to-well contamination can play a major role in microbiome studies, especially when using plate-based DNA extraction methods and for samples with low starting biomass. This type of contamination is difficult to detect and relatively infrequently discussed, but should be considered when designing and evaluating research. The majority of research to date has focused on identifying microbial contaminants in reagents and consumables [12,18,21] and subsequently using bioinformatics techniques to simply subtract out these contaminant taxa [22,27,32]. Existing tools to remove contaminant taxa or OTUs (operational-taxonomic units) from a dataset largely focus on these background contaminants, and don’t yet consider the case of contamination from proximal wells [27]. We show in this study that a large fraction of reads in the blank (NTC) samples originate from neighboring wells. In this study, we observed that contamination between samples can account for a significant fraction of the overall observed diversity in a sample, especially for no-template control blanks that are physically adjacent to relatively high-biomass samples. Given this, the simple approach of removing any taxa found in blanks is likely to remove the most prominent “real” taxa in a dataset. More sophisticated methods using additional information (such as the ‘decontam’ package [27]) are absolutely necessary in the face of well-to-well contamination, even for addressing the problem of reagent contaminants.

Identifying and removing well-to-well contamination *in silico* is challenging, as contamination events between wells are largely independent, and thus cannot be statistically identified and removed across a study in the same way that reagent contaminants are. However, several observations from this experiment should help researchers in planning experiments to minimize its effects. First, plate-based DNA extractions are much more susceptible to well-to-well contamination than the more painstaking tube-based extractions; for critical experiments, automated plate-based extractions should be carefully reconsidered. Second, even for tube-based extractions, well-to-well contamination was greatest in wells immediately adjacent to the source. Thus, sample location on plates should be explicitly considered in experimental design. When plating samples for extraction, it is important to block and/or randomize treatments across 96-well plates. Third, well-to-well contamination has the greatest impact in low-biomass samples, especially when they are processed adjacent to high-biomass samples that can act as sources. Because of this, it is important to have an awareness of the absolute concentration of microbial cells in samples, and to ensure that only samples of similar biomass are processed together. Lastly, when analyzing datasets, it is important to be aware that different methods will have different sensitivities to well-to-well contamination. For example, alpha-diversity estimates can be highly inflated by well-to-well contamination in samples with low starting diversity; and for beta-diversity estimates, binary metrics such as Jaccard or unweighted UniFrac are more likely to be affected than abundance-weighted metrics. Other experimental approaches to reduce the impacts of well-to-well contamination bear further investigation. These might include use of higher-fidelity liquid handling approaches [33], or broader adoption of unique-per-sample positive control spike-ins to allow the direct observation and statistical disambiguation of cross-contamination [34]. Methods which rely on identifying and subtracting putative contaminants from datasets need to be used with extreme caution, particularly if the identified sequence variants are present in primary samples.

Understanding experimental noise is extremely important for improving and guiding microbiome research best practices [23,24]. Specifically, addressing ‘hot’ negative controls is one of the great challenges to genomics based research. Since well-to-well contamination is an important component of this, we emphasize that for any given experiment, it is critical to identify any kit-specific background contaminants in a lot to best accurately remove contaminant taxa. While we have good power to estimate frequency of well-to-well contamination in our assays, extrapolating the frequency of well-to-well contamination in assays from other labs and methods is still a challenge. This suggests that while we can generalize to well-to-well contamination being a widespread problem, we can’t generalize the quantities or specifics. Further, this argues for other labs spending the effort to do similar in-house tests to evaluate their own pipelines. To identify these background contaminants, we recommend using a variety of positive controls titrations both at the DNA extraction stage and PCR stage [11]. Companies which manufacture high-throughput DNA extraction will need to invest in research and development to reduce well-to-well contamination. Lastly, measuring and accounting for well-to-well contamination identification and reduction will be critical for diagnostic research going forward [35–40].

## Conclusions

Contamination is a serious impediment to reproducibility in any genomics study, particularly microbiome research. As emerging diagnostic tests for environmental health and human health become more mainstream, it will be crucial for these tests to address variability in microbiome signal due to well-to-well contamination. Our study identified and quantified a previously undetected source of contamination in microbiome studies. We show that intensity of well-to-well contamination varies per extraction method with plate-based methods and lower biomass samples having higher rates of contamination. Our findings demonstrate the importance for the community to accept standards to best monitor and quantify these sources of noise in a given study.

## Methods

### Sample collection and processing

A total of 17 bacterial isolates including *Brevibacterium sp*, *Corynebacterium stationis*, *Brachybacterium sp*, *Arthrobacter sp*, *Propionibacterium acnes*, *Bacillus sp*, *Staphylococcus equorium*, *Staphylococcus succinus*, *Streptococcus angiosis*, *Desulfovibrio sulfodismutans*, *Serratia sp*, *Halomonas sp*, *Psychrobacter sp*, *Pseudomonas fragi*, *Vibrio rumo*, *Eschericia coli*, and *Vibrio fischeri* were collected and stored in PBS solution. The optical density, OD600, was measured for all isolates and the corresponding cell density estimated. Sixteen of these microbes (all except *V. fischeri*) were diluted to a final density of 1e8 cells per ml in a single 50 ml conical vial and were designated as ‘source’ organisms. The *V. fischeri* isolate was diluted to 1e6 cells per ml, designated as the ‘sink’ microbe, and stored in a single 50 ml conical. Both source and sink microbes were stored in a −80°C freezer until making aliquots for extractions. In addition, a mock community was created using these isolates by combining equal volume of all samples which also served as a reference for accounting for processing biases. An additional isolate of *Clostridium sp.* was measured and aliquoted into 16 different 2 ml tubes to be used for barcode testing. For DNA extraction at UCSD, 100 ul of ‘source’ and ‘sink’ samples were aliquoted into 2 96-well DNA extraction robot plates and 96 2-ml bead beating extraction tubes as indicated in the diagram (Additional file 1, Figure 1a). Following the Earth Microbiome Project protocol [2], the Qiagen PowerMag kit (Qiagen, Cat# 27500-4-EP) was used for robot extractions while the Qiagen DNeasy PowerSoil kit (Qiagen, Cat# 12888-100) was used for ‘manual, single-tube’ extractions. To test the effect of antifoam on reducing well-to-well contamination, we added 2 ul of antifoam-A concentrate (Sigma-Aldrich, Cat#A5633-25G) to half of each of the robot plates (Figure 1b-c). In addition to processing samples at UCSD, an additional 192 samples were plated (96) in a 96-well plate and 96 individual 2-ml bead beating tubes and sent to Argonne National lab in the same platemap scheme. The manual tube samples were processed using the Qiagen DNeasy PowerSoil kit (Qiagen, Cat# 12888-100) while the manual plate samples were processed using the Qiagen DNeasy PowerSoil HTP 96 kit (Qiagen, Cat# 12955-4).

### Amplicon sequencing

To distinguish between well-to-well contamination derived from DNA extraction versus PCR setup, each UCSD processed DNA extraction plate (2 robot plates and 1 manual plate) were subjected to two separate triplicate PCR reactions (Figure 1b-d). The mock community dilution plate and barcode testing plate were processed with a single triplicate PCR reaction each. The EMP 16S rRNA V4 primers 515f/806rB were used to amplify the samples. Equal concentrations of amplicons from each sample from all 8 plates were pooled and sequenced using a MiSeq [5,13,14]. The 192 samples DNA extracted at Argonne were processed using the same EMP primers and method but on a separate MiSeq run. Amplicon data was uploaded to Qiita [41] and processed with Qiime 1.9.1 [42]. Exact sequence tags from the first read were generated using the deblur pipeline under default parameters as described in the publication [43].

### Statistical analysis

Sequences processed with deblur were positively filtered against the reference database as part of the default workflow in deblur. In addition, singleton sequences were omitted from the dataset. The dataset was not rarified in order to best quantify well-to-well contamination for all samples processed. The sequence tags were identified for all of the positive controls used in this study and included in supplement (Additional Table 2). Sequences which did not have 100% match to those original controls were considered ‘background contaminants’ whereas the *Vibrio fischeri* deemed as ‘sink microbes’ and the 16 unique isolates deemed collectively as ‘source microbes’. For each of the 16 source microbes, 1 sink microbe, and 1 barcode leakage microbe, a custom script was used to generate 96-well plate maps to visualize well-to-well contamination. The distances of microbial dispersal ‘jumping’ was then calculated for each individual isolate using a custom script. Summary statistics of read counts, richness, and contamination metrics are summarized (Additional Table 3). To determine if well-to-well contamination was higher in robot compared to manual extractions, the composition of well-to-well contaminants was compared within NTCs, sink, and source independently, using the Kruskal-Wallis test. Further, to determine if well-to-well contamination was associated or more frequent with lower biomass samples, well-to-well composition was compared across the NTCs, sink, and source within each extraction method independently using Kruskal-Wallis test.

To determine the impact of well-to-well contamination on beta-diversity microbiome analyses, we calculated distance metrics of both Bray-Curtis [44,45] and Jaccard [46] and compared within categories. The three different extraction plates each had two separate PCR plates processed. The pairwise distances of unique Well_ID was calculated using both metrics for each of the two PCR plates belonging to each of the three DNA extraction plates. Sample types were grouped into NTCs (non-template control or blank), sink, or source. Within each group, the distances were compared using the Mann-Whitney test. To calculate effects for the entire pipeline which includes both PCR and DNA extraction, we combined the pairwise distances of the Well_IDs for each of the three DNA extraction plates (robot 1, robot 2, and manual) and grouped by sample type (NTC, sink, or source). Again, we compared the total dissimilarities of Bray-Curtis vs. Jaccard for each sample type using Mann-Whitney test.

## List of abbreviations

EMP: Earth Microbiome Project
NTC: non-template control (sterile water blanks)
sOTU: sub-operational taxonomic unit
sink: lower biomass microbial isolate used in experiment (approximately 100,000 cells)
source: higher biomass microbial isolate (1 of 16) (approximately 10,000,000 cells) Well_ID: refers to the well position (in a 96-well plate: A1-A12 → H1-H12) from which the sample was processed

## Acknowledgements

Thanks to Argonne National lab for processing samples.

## Data availability and materials

All data is made publically available on (Qiita ID 10401) and will be uploaded to public database ENA upon acceptance.

## Author contributions

JJM, AA, JS, GH, RK designed experiment

JJM, GH, JG processed samples

JJM, AA, JS analyzed and interpreted results

AA, JS developed custom scripts for various platemap analyses and distance-k

JJM, AA, JS, RK helped to write the manuscript

## Ethics approval

Not applicable

## Consent for publication

Not applicable

## Additional files

Supplemental Table S1 (full characterization of w2w and metadata)

Supplemental Table S2. Summary statistics on well-to-well contamination across DNA

Supplemental Figure S 1: Platemap descriptions of experimental design

Supplemental Figure S 2: The use of antifoam (antifoam = 1) does not reduce well-to-well contamination

Supplemental Figure S 3: Sources of contamination (well-to-well and background contaminants) across manual and robot extraction plate and PCR replicate plates. Summary of compositionality of NTCs (n-48) vs. sink (n=32) vs. source microbes (n=16) processed in two facilities across five DNA extraction plates (a) UCSD tube extraction, (b) UCSD plate extraction 1, (c) UCSD plate extraction 2, (d) Argonne tube extraction, (e) Argonne plate extraction. UCSD DNA extractions were processed each twice thus had two PCR per plate (PCR A, PCR B).

Supplemental Figure S 4: Summary composition of reads (median, inter-quartile range) of specific sample types: NTCs, sink, or source microbes.

Supplemental Figure S 5. Determining the origin of well-to-well contamination and its impact on distance metrics from 96 unique WellIDs across three DNA extraction plates and six PCR plates. (a) Summary comparison of use of compositional (Bray-Curtis) or presence-absence (binary Jaccard) to describe microbial communities from NTCs (red), sink microbes (lower biomass), or source microbes (higher biomass). (b) Determining the effects of well-to-well contamination from PCR processing only (PCR replicates) compared to the entire process of DNA extraction and PCR (DNA extraction replicates). The statistical tests are performed on dark colors only while lightly shaded bars indicate the replicates for robot extraction plates.

